# The N-linker region of hERG1a upregulates hERG1b potassium channels

**DOI:** 10.1101/2021.12.20.473560

**Authors:** Ashley A. Johnson, Taylor R. Crawford, Matthew C. Trudeau

## Abstract

A major physiological role of hERG1 (human Ether-á-go-go-Related Gene) potassium channels is to repolarize cardiac action potentials. Two isoforms, hERG1a and hERG1b, associate to form the native cardiac I_Kr_ current *in vivo*. Inherited mutations in hERG1a or hERG1b cause prolonged cardiac repolarization, Long QT Syndrome and sudden death arrhythmia. hERG1a subunits assemble with and enhance the number of hERG1b subunits at the plasma membrane, but the mechanism for the increase in hERG1b by hERG1a is not well understood. Here, we report that the hERG1a N-terminal PAS (Per-Arnt-Sim) domain-N-linker region expressed *in trans* with hERG1b markedly increased hERG1b currents and increased biotin-labelled hERG1b protein at the membrane surface. hERG1b channels with a deletion of the 1b domain did not have a measurable increase in current or biotinylated protein when co-expressed with hERG1a PAS domain-N-linker regions indicating that the 1b domain was required for the increase in hERG1b. Using a biochemical pull-down interaction assay and a FRET hybridization experiment, we detected a direct interaction between the hERG1a PAS domain-N-linker region and the hERG1b N-terminal 1b domain. Using engineered deletions and alanine mutagenesis, we identified a short span of amino acids at positions 216-220 within the hERG1a N-linker region that were necessary for the upregulation of hERG1b. Taken together, we propose that direct structural interactions between the hERG1a N-linker region and the hERG1b N-terminal 1b domain increase hERG1b at the plasma membrane. Mechanisms that enhance hERG1b current would be anticipated to shorten action potentials, which could be anti-arrhythmic, and may point toward hERG1b or the hERG1a N-linker as molecular targets for therapy for Long QT syndrome.

**Significance Statement:** Potassium (K) ion channels help to control the membrane voltage in cardiac cells and are responsible for maintaining the normal rhythm of the heart. A decrease in the number of K channels causes cardiac arrhythmia and sudden death. One K channel (KCNH2b, hERG1b) weakly forms channels at the cell membrane. Here we describe a new mechanism for upregulation of hERG1b. An N-terminal domain of a related K channel (KCNH2a, hERG1a), expressed *in trans* as a separate piece, interacts with hERG1b and increases the number of hERG1b channels at the membrane. A loss of hERG channels is pro-arrhythmic and an increase in hERG channels is considered anti-arrhythmic, thus the hERG1a N-terminal domain is a potential therapeutic for cardiac arrhythmias.

## Introduction

The human Ether-á-go-go-Related Gene (hERG, KCNH2) encodes a voltage-activated potassium channel that is critical in human health and disease. Two isoforms of the hERG gene, hERG1a and hERG1b, form the rapid component of the delayed rectifier potassium current (I_Kr_) in heart (1-5) which repolarizes the ventricular action potential (6). Mutations in hERG1a (7) or hERG1b (8) are linked to Long QT Syndrome (LQTS), a predisposition to prolonged action potentials, cardiac arrhythmia, and sudden death. hERG channels are the targets for acquired LQTS which is due to the off-target inhibition of hERG channels by drugs (5) and is a common clinical problem (9).

hERG1a and hERG1b isoforms have different structural and functional properties that are due in part to their divergent N-terminal regions. The hERG1a N-terminal region has 396 amino acids whereas the N-terminal region of hERG1b has 59 amino acids (Fig. 1A, B). The N-terminal region of hERG1a contains a PAS (Per-Arnt-Sim) domain and PAS-CAP, which together encode the first 135 amino acids of hERG1a (Fig. 1A). The first 135 amino acids of hERG1a were initially termed the ‘eag domain’ (10), but here we will follow the recent suggestion that the first 135 amino acids be referred to as the PAS domain for clarity (11). In hERG1a, the PAS domain is followed by an N-linker region, encoded by amino acids 136-396, which connects the PAS domain and the S1 transmembrane domain (Fig. 1A). The function of the N-linker region is not well understood. In contrast to mammalian ERG1a, mammalian ERG1b does not contain a PAS domain or N-linker region and instead the first 36 amino acids of hERG1b are unique (3) and not conserved with other proteins (Fig. 1B). hERG1a and hERG1b are identical from N-terminal residues 377 in hERG1a and 37 in hERG1b through the carboxyl terminus (Fig. 1A, B). hERG1a channels are characterized by a slow time course of deactivation (closing) that requires the PAS domain (10, 12, 13) and the C-terminal CNBHD (Cyclic Nucleotide Binding Homology Domain) (12, 13). The mechanism for slow deactivation requires a direct structural interaction between the PAS and CNBHD domain in an intersubunit, domain-swapped arrangement (12, 13) that is validated by a CryoEM structure of the hERG1a channel (14). The PAS-CNBHD interaction is conserved in other KCNH (KCNH1, EAG) channels (15) and may be a defining feature of KCNHs. Mouse and human ERG1b channels have rapid deactivation (3, 16) that is due to the lack a PAS domain (16). Slow deactivation can be introduced in hERG1b channels by co-expression *in trans* with hERG1a PAS domains (16), indicating that in hERG1a/hERG1b heteromeric channels the hERG1b CNBHD likely makes an interaction with the PAS domain of hERG1a (16).

**Figure 1.**
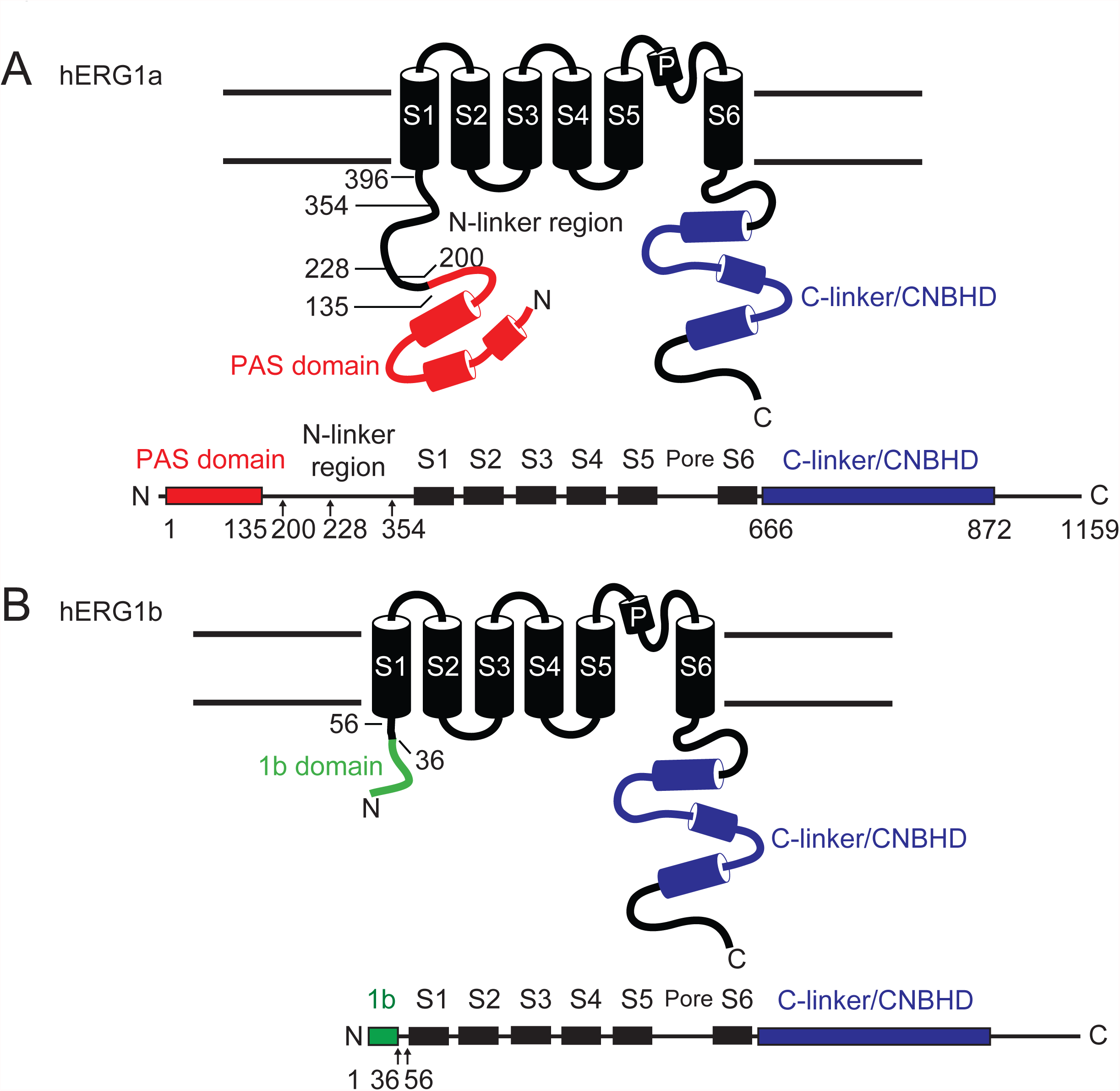
Scheme of hERG1a and hERG1b subunits. hERG1a and hERG1b contain 6 transmembrane domains and C-terminal C-linker and CNBHDs. hERG1a and hERG1b diverge in the N-terminal regions. A) The N-terminal of hERG1a is comprised of 396 amino acids and contains a PAS domain and N-linker region. B) The hERG1b N-terminal lacks a PAS domain and N-linker region and instead contains 56 amino acids, the first 36 of which are unique and referred to as the 1b domain.

hERG1a subunits form homomeric channels with large currents in heterologous expression systems (4, 5, 17). In contrast, hERG1b (and mouse ERG1b) can form homomeric channels but the currents are very small, as hERG1b is preferentially retained at intracellular membranes due in part to a di-arginine retention signal in its unique N-terminal domain (3, 16, 18, 19). Mammalian ERG1a and ERG1b subunits directly associate in heterologous and native cells to form hetero-tetrameric channels (1-3, 8, 18-22). Co-expression of hERG1a with hERG1b increases the biochemical maturation of hERG1b subunits (19), suggesting that hERG1a helps to increase hERG1b at the plasma membrane. A biochemical interaction was detected between fusion proteins encoding the N-terminal region-S2 transmembrane domains of hERG1a and N-terminal region-S2 of hERG1b (18), but the functional correlate of this interaction is not clear and more precise molecular determinants for the N-termini S2 interaction have not been determined.

In this study, we use hERG1a N-terminals composed of the PAS domain and N-linker regions (and bearing deletions or mutations) encoded by mRNAs in *Xenopus* oocytes or encoded by cDNAs in HEK293 cells and co-expressed *in trans* with hERG1b channels to simultaneously probe functional and structural interactions. Using hERG1a N-terminals expressed *in trans* with hERG1b channels has several advantages as an experimental approach: 1) Changes in hERG1b channel properties can be attributed directly to the hERG1a N-terminals, providing a direct link between structure of the hERG1a N-terminals and hERG1b channel function, and 2) changes in hERG1b channel properties are not obscured by the ion conducting properties of intact hERG1a channels, as in previous studies and 3) since the hERG1a PAS domain regulates hERG1b channel gating properties, regulated gating is an independent internal positive control for expression of hERG1a PAS-domain-N-linkers.

Here, we report that hERG1a N-terminals encoding the PAS domain and N-linker region increased hERG1b, as measured by an increase in hERG1b ionic current recorded with two-electrode voltage-clamp or whole-cell patch-clamp, and by an increase in the amount of hERG1b at the plasma membrane as measured with surface biotinylation and Western blot assays. hERG1b channels with a deletion of the unique N-terminal 1b domain did not have a measurable increase in current or biotinylated protein when co-expressed with hERG1a PAS domain-N-linker regions, indicating that the 1b domain was required for the increase in hERG1b. Using biochemical pull-down interaction assays and FRET two-hybrid interaction assays, we detected an interaction between the hERG1a PAS-domain-N-linker region and the hERG1b N-terminal 1b domain. The hERG1a PAS domain-N-linker region-dependent increase of hERG1b required the hERG1a N-linker, since hERG1a PAS domains expressed *in trans* with hERG1b did not measurably increase hERG1b currents or biotinylated hERG1b protein. Using hERG1a PAS domain-N linker regions with sequential deletions within the N-linker, we determined that residues 200-228 of the N-linker were required for the increase in hERG1b. We used alanine mutagenesis to further examine the N-linker region bounded by amino acids 200 and 228 and determined that residues 216-220 in the hERG1a N-linker region were necessary for the increase in hERG1b. Mechanisms that enhance hERG1b current are anticipated to shorten action potentials, which could be anti-arrhythmic, and may point toward hERG1b or the hERG1a N-linker as molecular targets for therapy for Long QT syndrome.

## Results

To carry out these experiments, we expressed hERG1b in *Xenopus* oocytes and used two-electrode voltage-clamp to record hERG1b currents, which are characterized by rapid channel deactivation and relatively small ionic currents (Fig. 2A-E). We next co-expressed the hERG1a PAS domain-N-linker region fused to CFP (hERG1a N1-354-CFP) *in trans* with hERG1b fused to Citrine (Fig. 2A). Compared to wild-type hERG1b channels, we found that the hERG1a PAS domain-N-linker region regulated and slowed the deactivation time course of hERG1b (Fig. 2A, B) and shifted the steady-state activation relationship of hERG1b toward more negative voltages (Fig. 2C, Table 1). We also report here that the hERG1a PAS domain-N-linker region markedly increased the magnitude of hERG1b currents (Fig. 2A, D, E). We found similar results using whole-cell patch-clamp recordings in HEK293 cells, where a hERG1a PAS domain-N-linker region markedly increased the magnitude of hERG1b currents (Fig. S1). To directly test whether the hERG1a PAS domain-N-linker region increased the number of hERG1b channels at the plasma membrane, we measured surface biotinylated hERG1b using SDS-PAGE and Western blot analysis in HEK293 cells (Fig. 2F). Biotinylated hERG1b (Fig. 2F, lane 1) was detected as an upper band at 120 kDa and a lower band at 110 kDa on a Western blot (arrows, Fig. 2F). Mature (upper) hERG1b bands were faint as measured with densitometry (Fig. 2G), consistent with small hERG1b currents (Figs. 2A-E, S1). In marked contrast, the hERG1a PAS domain-N-linker region co-expressed *in trans* with hERG1b increased the intensity of the mature hERG1b band (Fig. 2F, lane 2; Fig. 2G). These biochemistry results are consistent with our electrophysiology results showing that the hERG1a PAS domain-N-linker region increased hERG1b currents (Figs. 2, S1).

**Table 1:** Properties of hERG channels expressed in Xenopus oocytes.

| Channel | Current<br>amplitude<br>(μA) | SEM | Time constant<br>of deactivation<br>(ms) | SEM | $V_{1/2}$ | $\kappa$ |
| --- | --- | --- | --- | --- | --- | --- |
| hERG1b | 0.67 | 0.11 | 0.03 | 0.00 | $-24.1 \pm 1.19$ | $10.944 \pm 1.07$ |
| hERG1b + hERG1aN1-135 | 0.86 | 0.03 | 0.23 | 0.05 | $-32.639 \pm 0.76$ | $9.3495 \pm 0.63$ |
| hERG1b + hERG1aN1-200 | 0.87 | 0.12 | 0.42 | 0.09 | $-36.643 \pm 0.59$ | $7.7255 \pm 0.59$ |
| hERG1b + hERG1aN1-228 | 2.42 | 0.44 | 0.31 | 0.04 | $-43.644 \pm 0.85$ | $8.537 \pm 0.80$ |
| hERG1b + hERG1aN1-354 | 2.27 | 0.49 | 0.33 | 0.01 | $-41.165 \pm 0.51$ | $7.7639 \pm 0.56$ |
| hERG1bΔ2-36 | 0.45 | 0.01 | 0.03 | 0.01 | $-25.586 \pm 4.15$ | $11.823 \pm 3.68$ |
| hERG1bΔ2-36 + hERG1aN1-354 | 0.85 | 0.09 | 0.29 | 0.03 | $-57.73 \pm 0.661$ | $7.9077 \pm 0.69$ |
| hERG1b + hERG1aN1-228<br>(TPAAP→AAAAA) | 2.25 | 0.17 | 0.33 | 0.02 | $-43.844 \pm 0.15$ | $7.4301 \pm 0.14$ |
| hERG1b + hERG1aN1-228<br>(SSESL→AAAAA) | 1.20 | 0.05 | 0.34 | 0.03 | $-46.403 \pm 0.51$ | $7.6938 \pm 0.40$ |
| hERG1b + hERG1aN1-228<br>(ALDEV→AAAAA) | 3.56 | 0.50 | 0.29 | 0.06 | $-44.762 \pm 0.91$ | $7.7667 \pm 0.81$ |
| hERG1b + hERG1aN1-228<br>(TAMDN→AAAAA) | 0.73 | 0.18 | 0.44 | 0.05 | $-41.812 \pm 0.31$ | $7.1849 \pm 0.36$ |
| hERG1b + hERG1aN1-228<br>(HVAGL→AAAAA) | 2.05 | 0.30 | 0.48 | 0.02 | $-42.842 \pm 0.72$ | $6.4448 \pm 0.83$ |

**Figure 2.**
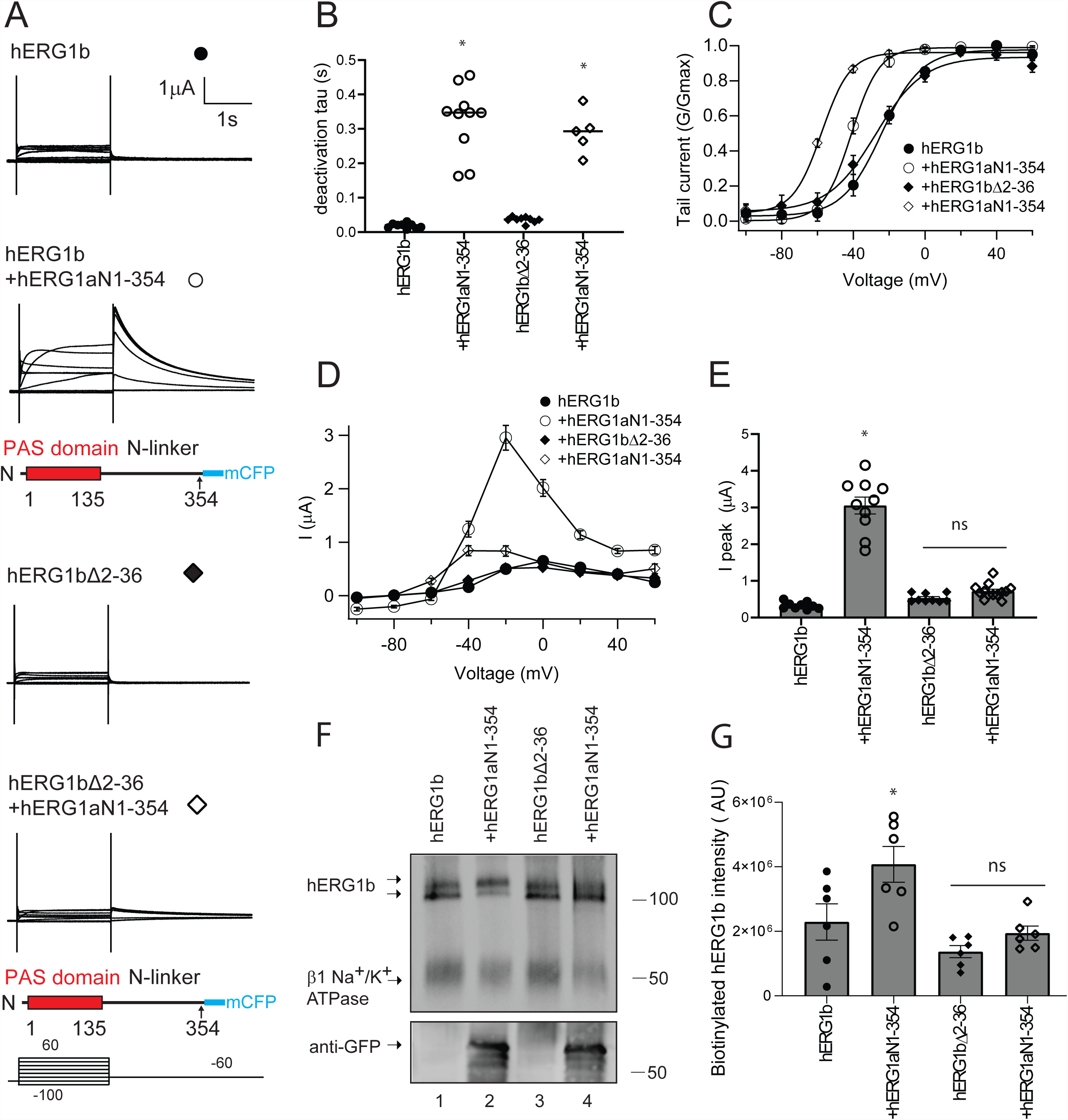
hERG1a PAS domain-N-linker regions increase hERG1b currents and surface expression but do not measurably increase currents or surface expression of hERG1bΔ2-36 channels in which the 1b domain is deleted. A) Two-electrode voltage-clamp recordings of hERG1b (closed circles), hERG1b expressed *in trans* with the hERG1a PAS-N-linker (open circles), hERG1b Δ2-36 (closed diamonds) and hERG1b Δ2-36 expressed *in trans* with the hERG1a PAS-N-linker (open diamonds). Voltage pulse protocol as indicated. Scale bar is 1 μA and 1 s. B) Time constant of deactivation measured at -60 mV. C) Conductance-voltage (G-V) relationship. The solid line is a Boltzmann fit to the data. D) Current voltage (I-V) relationship. E) Peak current measured at a depolarizing voltage command from data in A. F) Western blot of surface biotinylated hERG1b or hERG1b Δ2-36 expressed individually (lanes 1,3) or *in trans* with the hERG1a PAS-N-linker (lanes 2,4) blotted with the anti-hERG KA antibody. Na/K-ATPase served as a loading control. Anti-GFP antibody detects hERG1a PAS domain-N-linker regions fused to mCFP as an input control. G) Plot of densitometry of data from F. N ≥ 3 for each. Error bars are mean ± SEM.

To identify functional determinants in hERG1b that were necessary for the increase in hERG1b by the hERG1a PAS domain-N-linker region, we generated a hERG1b channel that lacked the unique hERG1b N-terminal 1b domain (hERG1b Δ2-36-Citrine). hERG1bΔ2-36 had rapid deactivation, a conductance-voltage plot midpoint of -25 ± 4 mV (Fig. 2A-C), small outward currents (Fig. 2 A, D, E) and a low density of mature protein at the plasma membrane (Fig. 2F, lane 3; G), properties similar to that in wild-type hERG1b (Figs. 2A-G, Table 1). When we co-expressed the hERG1a PAS domain-N-linker *in trans* with hERG1bΔ2-36 channels, we did not detect a measurable increase in ionic currents or biotinylated protein (Fig. 2A, D, E, F, lanes 3 and 4; G). These results suggest that the hERG1b N-terminal 1b domain was necessary for hERG1b to be increased by the hERG1a PAS domain-N-linker. In positive control experiments, the hERG1a PAS domain-N-linker slowed hERG1bΔ2-36 deactivation (Fig. 2 A, B), left-shifted the G-V relationship (Fig. 2C) and was detected on a Western blot (Fig. 2F, lane 4), which together are evidence that hERG1a PAS domain-N-linker region was indeed expressed in these experiments but did not measurably upregulate hERG1bΔ2-36 channels.

Next, we performed biochemical pull-down interaction assays to test for a direct interaction between the hERG1b N-terminal 1b domain and the hERG1a PAS domain-N-linker region (Fig. 3). An advantage of the pull-down technique is that two fusion proteins that interact in this assay are considered to be sufficient to make a direct protein-protein interaction. We generated the entire N-terminal domain of hERG1b as a glutathione-S-transferase (GST) fusion protein (GST-hERG1b N1-56) and the hERG1a PAS domain-N-linker region as a 6x-histidine fusion protein with a FLAG epitope for detection (6xHis-hERG1a N1-354-FLAG). We expressed each fusion protein in bacteria and purified them using GST beads or nickel columns, respectively. GST and GST-hERG1b N1-56 inputs are shown on the Coomassie Blue (CB) gel (Fig. 3, upper panel). We incubated GST or GST-hERG1b N1-56 with the 6xHis-hERG1a N1-354-FLAG and performed SDS-PAGE and Western blot analysis. We did not detect the hERG1a PAS domain-N-linker region in negative control experiments with GST (Fig. 3, lower panel, lane 1). In contrast, we detected a band corresponding to the 6xHis-hERG1a N1-354-FLAG at its predicted MW (37kD; Fig. 3, lower panel, lane 2) indicating that the hERG1a PAS domain-N-linker region made an interaction with the hERG1b N-terminal domain.

**Figure 3.**
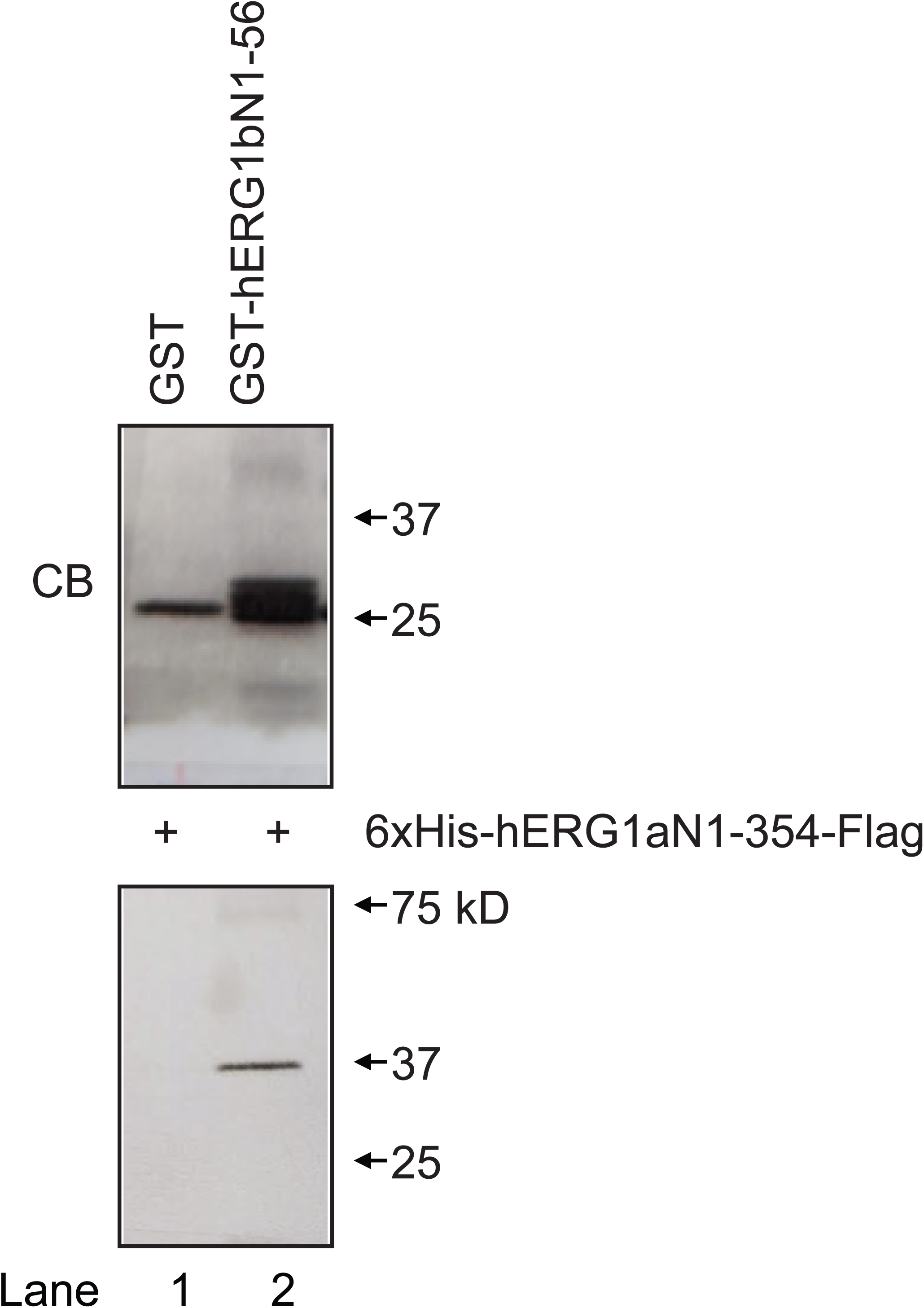
The hERG1b N-terminal domain interacts with the hERG1a PAS domain-N-linker region in a biochemical pull-down interaction assay. GST or GST– hERG1b N-terminal domains were incubated with hERG1a PAS domain-N-linker regions as indicated (+). Upper Panel) Coomassie blue (CB)-stained gel of GST and GST-hERG1b N-terminal domain (GST-hERG1b N1-56) inputs. Lower Panel) Western blot following GST (lane 1) or GST-hERG1b N-terminal domain (lane 2) incubation with the hERG1a PAS domain-N-linker region (hERG1a N1-354). The band at 37kD in lane 2 corresponds to the hERG1a PAS-N-linker and indicates a specific interaction. hERG1a PAS-N-linker fusion proteins were detected with an anti-Flag antibody.

Next, in an experiment which complements the biochemical pull-down interaction assay, we examined interactions between the hERG1b N-terminal domain and hERG1a intracellular domains using a FRET two-hybrid assay (Fig. 4). In a FRET two-hybrid experiment (23), two protein domains (or regions) of interest are each fused to different fluorescent proteins that are FRET pairs (we used CFP as the FRET donor and Citrine as the FRET acceptor) as in previous studies (12). The proteins domains fused to FRET pairs are co-transfected and allowed to freely associate in the milieu of a biological cuvette (e.g., a HEK293 cell). If FRET is detected in this experiment, it suggests that the two protein domains associated and brought the CFP and Citrine within sufficient proximity (within 80 Angstroms) for energy transfer (12, 23). We generated the hERG1b N-terminal domain fused to Citrine fluorescent protein (hERG1b N1-56-Citrine) and tested for FRET in an unbiased screen by co-expressing it with the hERG1a PAS domain-N-linker region (hERG1a N1-354-CFP) or hERG1 C-terminal region, which contains the C-linker and CNBHD (hERG1 C666-1159-CFP) (Fig. 4 A, B). The hERG1 C-terminal region is identical in hERG1a and hERG1b (Fig. 1). We co-expressed each FRET pair in HEK293 cells at similar ratios (Fig. 4D) and performed fluorescence imaging and ratiometric analysis of fluorescence spectra (Fig. 4A, B) for each combination of donors and acceptors. We calculated Ratio A – Ratio A_0_, which is a value proportional to FRET efficiency, where a value greater than zero indicates FRET (Fig. 4C) (see Methods). We detected robust FRET between the hERG1b N-terminal domain (hERG1b 1-56-Citrine) and the hERG1a PAS domain-N-linker region (hERG1a N1-354-CFP) (Fig. 4A, C). These results suggest that the hERG1a PAS-domain-N-linker makes an interaction with the hERG1b N-terminal domain and directly supports our similar results and conclusion from the biochemical pull-down interactions (Fig. 3) and electrophysiology and biotinylation experiments (Fig. 2). We did not detect measurable FRET between the hERG1b N-terminal domain and the hERG1 C-terminal domain (hERG1 C-666-1159-CFP) (Fig 4. B, C). These results show that the interaction of the hERG1b N-terminal domain with the hERG1a PAS domain-N-linker region was specific.

**Figure 4.**
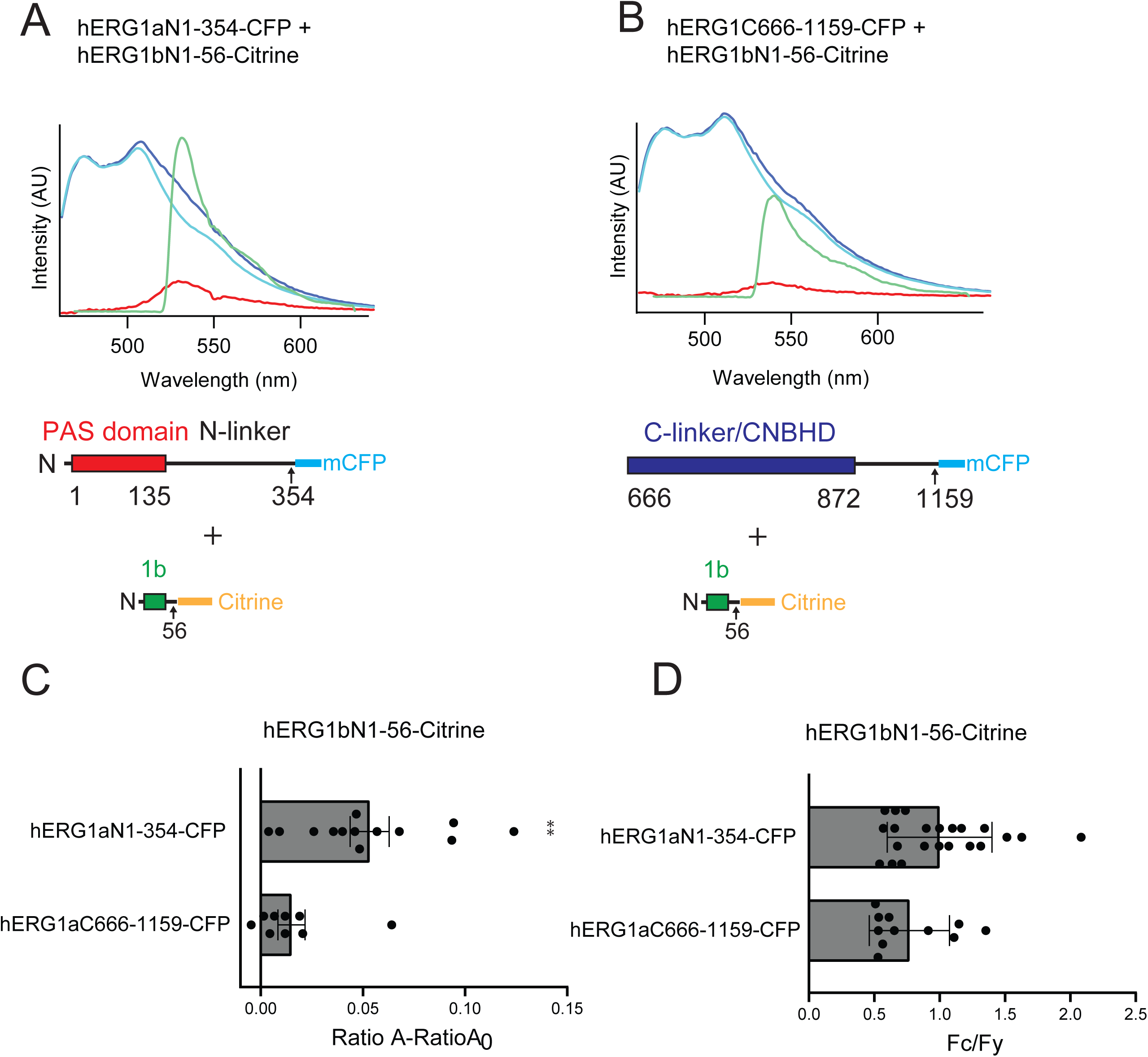
The hERG1b N-terminal domain and the hERG1a PAS domain-N-linker region associate in a FRET two-hybrid assay. Fluorescence intensity (A.U) vs. wavelength (nm) for A) the hERG1a PAS domain-N-linker region-mCFP and hERG1b N-terminal domain-Citrine and B) the hERG1 terminal region-mCFP and hERG1b N-terminal domain-Citrine. Traces are emission spectrum for CFP control (cyan), Citrine (green), donor (CFP) and acceptor (Citrine; blue trace) and extracted spectrum (red, see Methods). C) Plot of RatioA - RatioA_0_ (a value proportional to FRET) for each pair of hERG proteins, as indicated. D) Plot of peak CFP fluorescence intensity (Fc) divided by peak Citrine fluorescence intensity (Fy), as a control for similar amounts of input. Error bars are mean ± SEM.

To identify functional and structural determinants within the hERG1a PAS domain-N-linker that were required for the increase in hERG1b, we next co-expressed the hERG1a PAS domain (without the N-linker region) fused to CFP (hERG1a N1-135-CFP) *in trans* with hERG1b (Fig. 5A). The resulting currents (Fig. 5A) had a slower time course of deactivation (Fig. 5B) and a slightly left-shifted G-V (Fig. 5C, Table 1) when compared to hERG1b, as anticipated (16). However, we did not detect a marked change in magnitude of outward current during depolarization as compared to hERG1b (Fig. 5A, D, E). In a side-by-side control experiment, the hERG1a PAS domain-N-linker increased hERG1b current (Fig. 5A, D, E) as in Figure 2. When the hERG1a PAS domain was co-expressed *in trans* with hERG1b, we did not detect a measurable change in density of hERG1b bands (Fig. 5F, lane 2; G), consistent with the lack of upregulation of hERG1b by the hERG1a PAS domain in electrophysiology experiments (Figs. 5A, D, E; S1). In a side-by-side control experiment, the hERG1a PAS domain-N-linker increased hERG1b maturation (Fig. 5F, lane 3; G). Together, our results confirm that the hERG1a PAS domain regulates gating in hERG1b, as previously reported (16, 24), and we show that the hERG1a N-linker region was necessary to increase the magnitude of hERG1b currents and to increase the amount of hERG1b channels at the plasma membrane.

**Figure 5.**
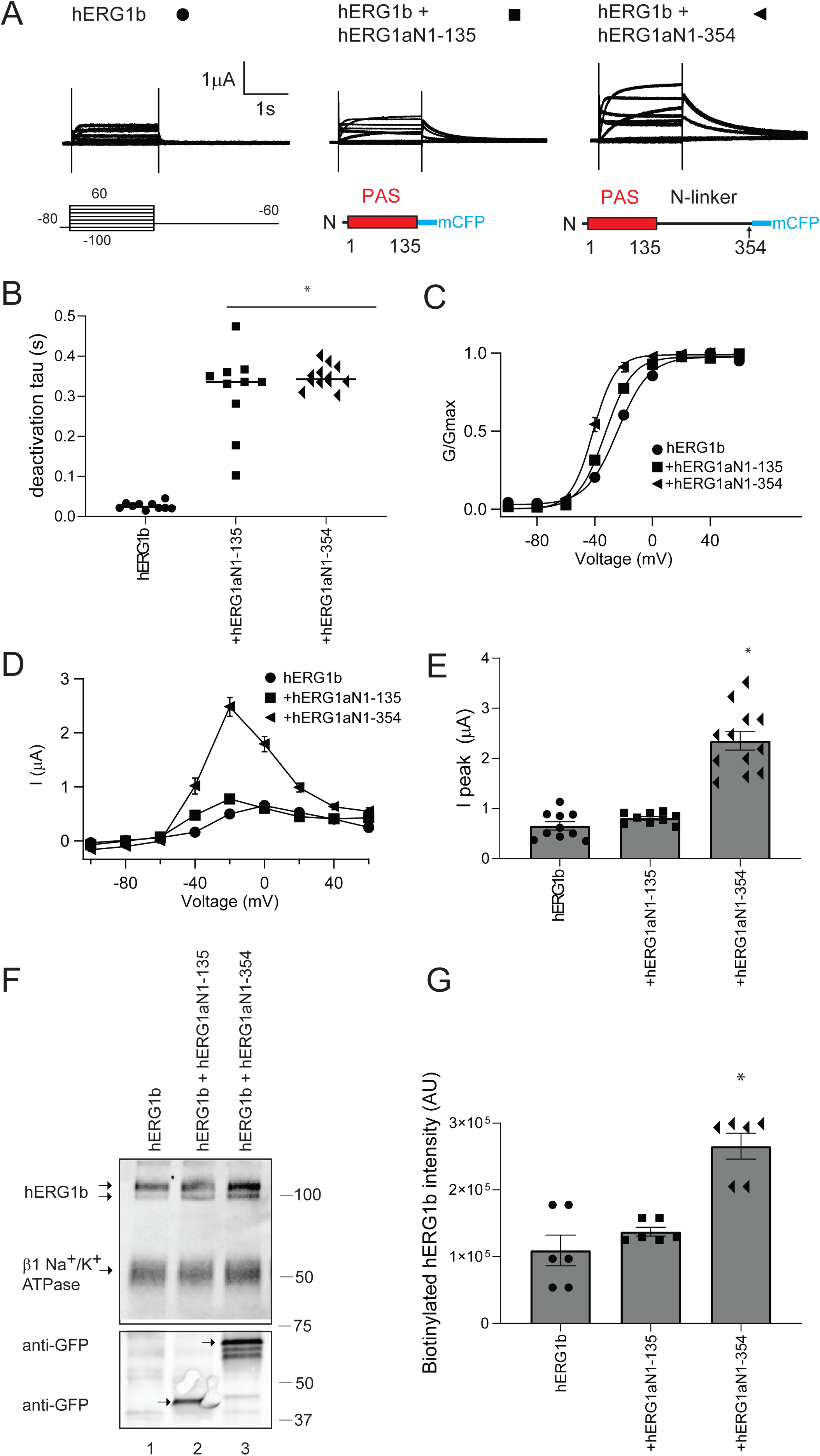
The hERG1a PAS domain did not increase hERG1b current. A) Two-electrode voltage-clamp recordings of hERG1b, hERG1b expressed *in trans* with the hERG1a PAS domain and hERG1b expressed *in trans* with the hERG1a PAS domain-N-linker. Voltage protocol is the same as in Fig. 2. Scale bar is 1 μA and 1 s. B) Plot of time constant of deactivation at -60 mV. C) Conductance-voltage (G-V) plot D) Current-voltage (I-V) plot. E) Histogram of peak current amplitude at a depolarizing voltage from data in A. N ≥ 9 for each. Error bars are mean ± SEM. F) Western blot of surface biotinylated hERG1b co-expressed with control vector (lane 1), hERG1a PAS domain (lane 2) and hERG1a PAS domain-N-linker region (lane 3) blotted with the anti-hERG KA antibody. Na^+^/K^+^ ATPase was the loading control. Anti-GFP antibody detects hERG1a PAS domain and PAS domain-N-linker region proteins as an input control. G) Histogram of band intensity of biotinylated hERG1b. N ≥ 3 for each. Error bars are mean ± SEM.

Our results suggested that a part of the hERG1a N-linker region located between amino acids 135 and 354 was necessary for the functional increase in hERG1b channels at the membrane (Fig. 6). To identify a region between residues 135 and 354 that was required to increase hERG1b, we generated constructs that encoded the hERG1a PAS domain and up to amino acid 200 of the N-linker region (hERG1a N1-200) and another that encoded the PAS domain and up to amino acid 228 of the N-linker region (hERG1a N1-228) (Fig. 6A). When co-expressed *in trans*, we found that hERG1a N1-200 regulated hERG1b gating but did not enhance outward currents or biotinylated hERG1b (similar to hERG1a N1-135) whereas hERG1a N1-228 regulated gating and also increased hERG1b currents and biotinylated hERG1b at the membrane (similar to hERG1a N1-354) (Fig. 6A-G). These results suggested that a region between amino acid 200 and 228 in the hERG1a N-linker was necessary for the increase in hERG1b.

**Figure 6.**
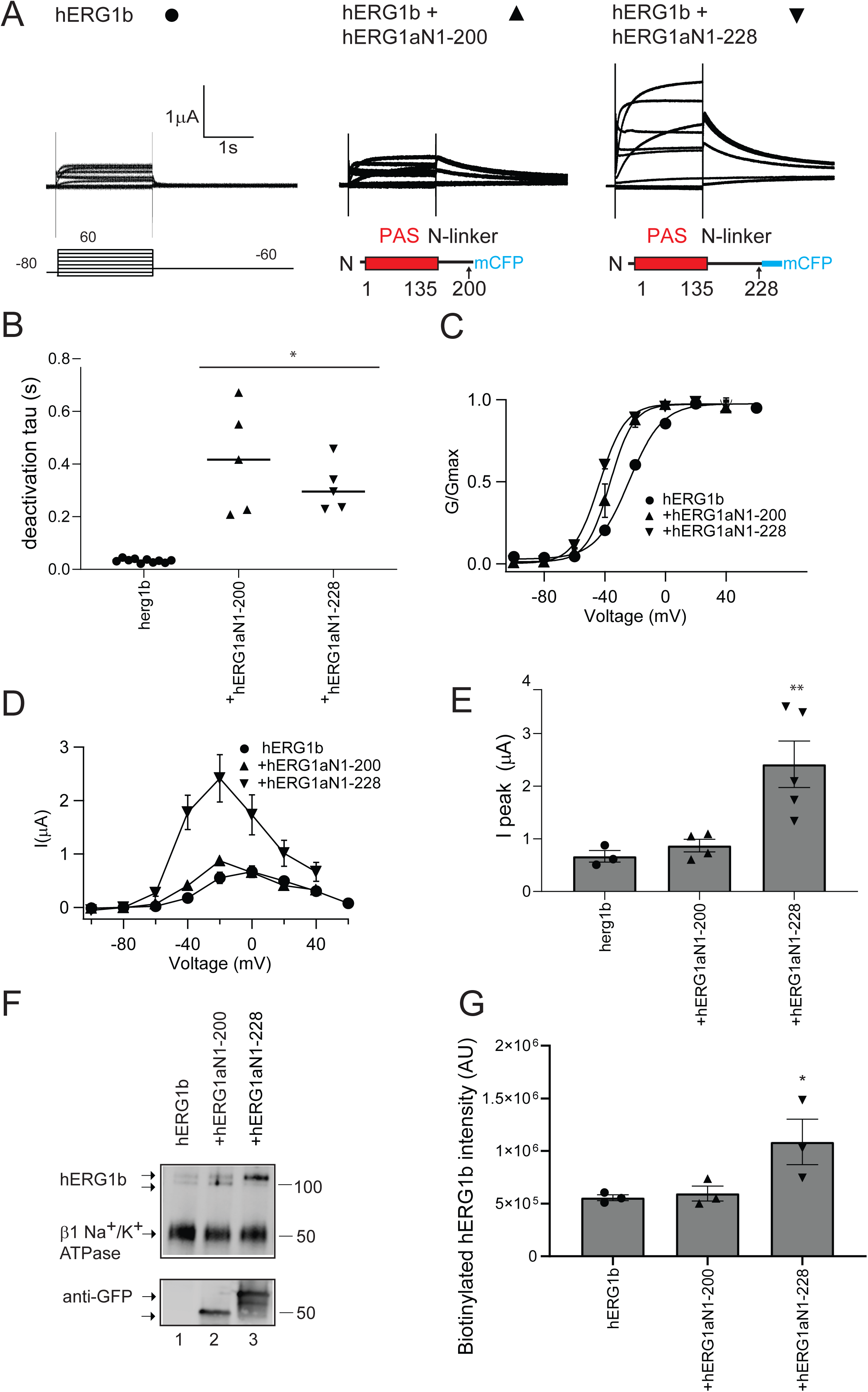
hERG1a PAS domain-N-linker regions encoding amino acids 1-228, but not 1-200, increased hERG1b current. A) Two-electrode voltage-clamp recordings of hERG1b and hERG1b co-expressed *in trans* with hERG1a PAS domain-N-linker regions, in which the N-linker is truncated at amino acid 200 or amino acid 228, as indicated. Voltage protocol is the same as in Figure 2. Scale bars are 1 μA and 1 s. B) Plot of time constant of deactivation at -60 mV. C) Conductance-voltage (G-V) plot. D) Current-voltage (I-V) plot. E) Histogram of peak current amplitude at a depolarizing voltage from data in A. F) Western blot of surface biotinylated hERG1b co-expressed *in trans* with control vector (lane 1), hERG1a residues 1-200 (lane 2) or hERG1a residues 1-228 (lane 3) and detected with anti-hERG KA. Na^+^/K^+^ ATPase was the loading control. Anti-GFP antibody detects hERG1a 1-220 or 1-228 as input controls. G) Plot of densitometry (A.U.) of hERG1b from data as in F. N ≥ 3 for each. Error bars are mean ± SEM.

To identify amino acids between hERG1a residues 200 and 228 required to increase hERG1b, we used alanine mutagenesis. We generated five mutants each with 5 consecutive alanine substitutions spanning the region from residues 200 to 228 in the background of hERG1a N1-228 (Fig. 7A). We co-expressed each of these, e.g., hERG1a N1-228 (^201^TPAAP^205^ to ^201^AAAAA^205^) *in trans* with hERG1b and performed two-electrode voltage-clamp recordings (Fig. 7B), whole-cell patch-clamp recordings (Fig. S1), and biotinylation experiments (Fig. 8). We found that each of the five alanine mutants regulated hERG1b deactivation kinetics (Fig. 7B, C) and were detected on Western blots (Fig. 8) indicating that they were all expressed. Of these, four of the mutants (^201^TPAAP^205, 206^SSESL^210, 211^ALDEV^215, 221^HVAGL^226^) increased the hERG1b current amplitude (Fig. 7 B, E, F) and increased biotinylated hERG1b at the membrane (Fig. 8A-C, E). In contrast, the mutant ^216^TAMDN^220^ neither increased the hERG1b current (Fig. 7B, E, F; Fig. S1) nor increased biotinylated hERG1b at the membrane (Fig. 8 D). We conclude from these data that the residues ^216^TAMDN^220^ within the hERG1a N-linker were required for the increase in hERG1b current. These findings represent a new role for the hERG1a N-linker in upregulating hERG1b.

**Figure 7.**
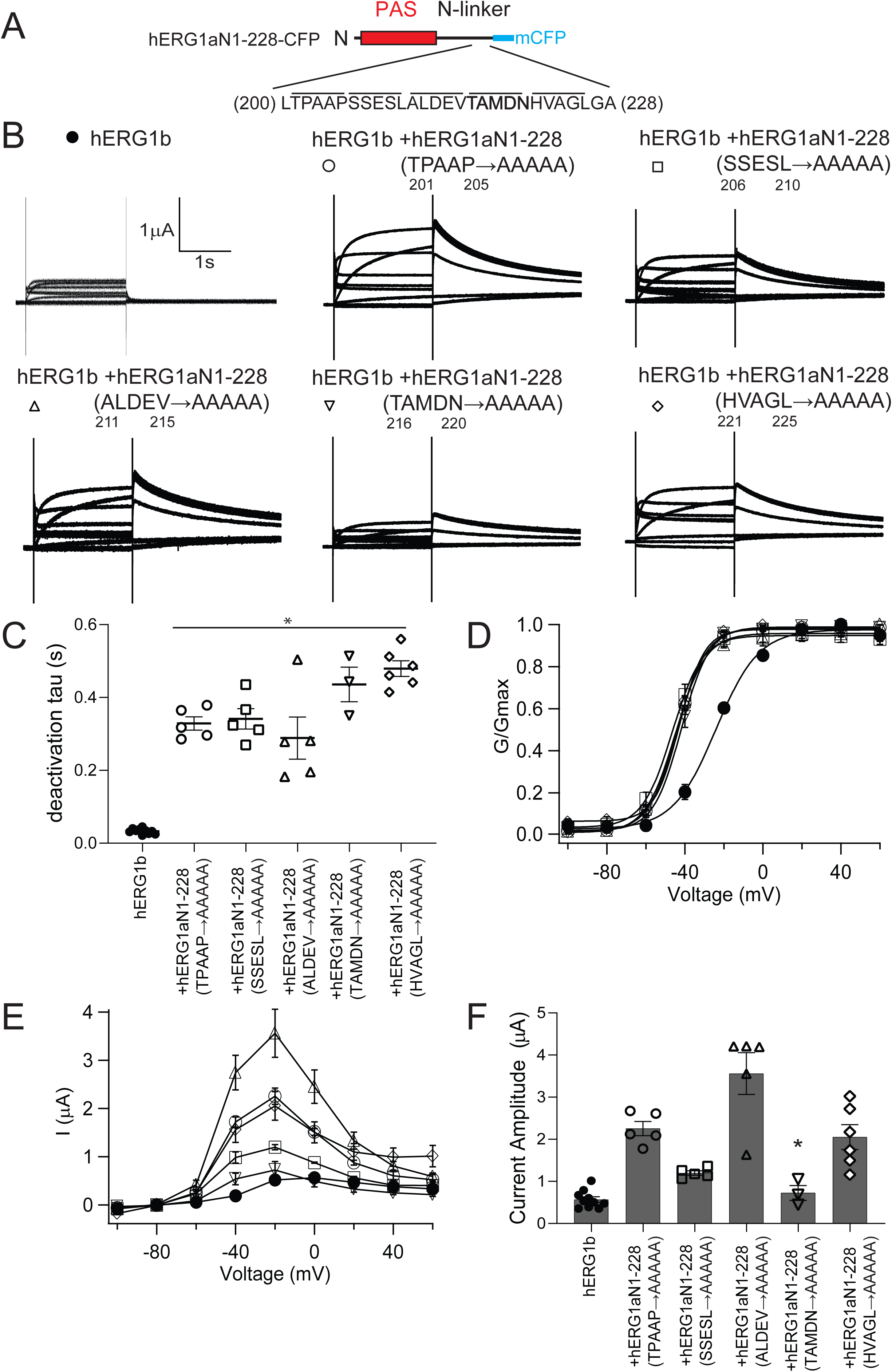
Alanine mutagenesis revealed amino acids 216-220 of the hERG1a N-linker were necessary for increased hERG1b current amplitude. A) Amino acid sequence of hERG1a residues 200-228 with each line denoting a group of 5 sequential residues that were mutated to alanine. B) Exemplar two-electrode voltage-clamp recordings of hERG1b and hERG1b co-expressed *in trans* with each hERG1a PAS domain-N-linker containing the alanine mutants as labeled and as depicted in panel A. C) Plot of time constant of deactivation at -60 mV. D) Conductance-voltage (G-V) plot. E) Current-voltage (I-V) plot. F) Histogram of peak current amplitude at depolarizing voltage from data in A. N ≥ 3 for each. Error bars are mean ± SEM.

**Figure 8.**
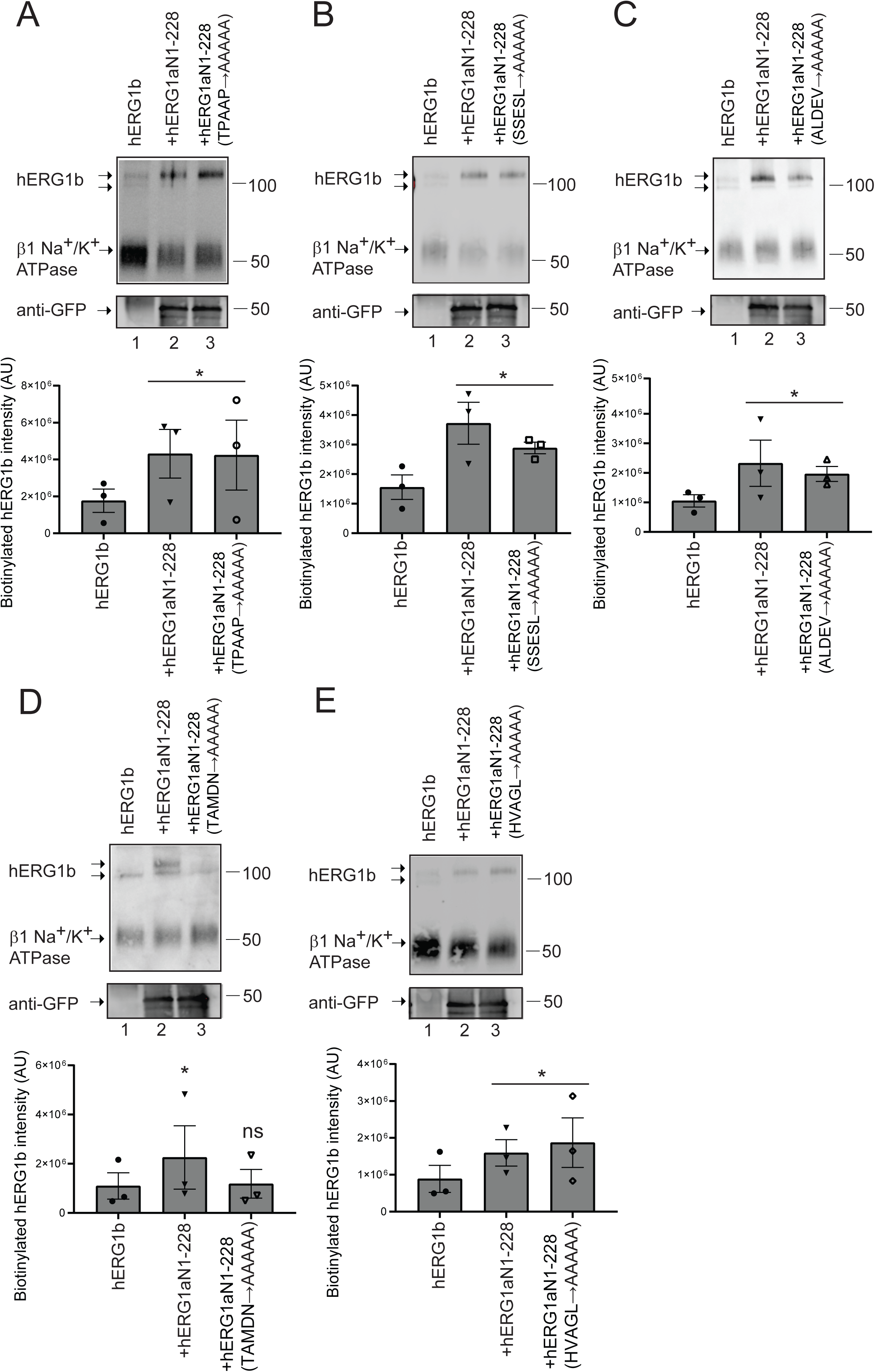
Alanine mutagenesis revealed amino acids 216-220 of the hERG1a N-linker were necessary for increased hERG1b maturation and cell surface expression. A-E) Western blot of surface biotinylated hERG1b co-expressed *in trans* with control vector (lane 1), hERG1a N1-228 (lane 2) or a hERG1a N1-228 bearing 5 alanine mutants as labelled (lane 3) and as depicted in panel A of Fig. 8 and blotted with anti-hERG KA. Na^+^/K^+^ ATPase was the loading control. Anti-GFP antibody was used to detect hERG1a N1-228 and each N1-228 alanine mutant as an input control. Plot of densitometry (A.U.) of data from Western blots. N = 3 for each. Error bars are mean ± SEM.

## Discussion

To explain our data, we propose that the hERG1a N-terminal PAS domain-N-linker region increases hERG1b current by interacting with and stabilizing (large arrow) hERG1b channels at the plasma membrane (Fig. 9A). In contrast, hERG1b alone only weakly makes homomeric channels at the plasma membrane (small arrow) and instead is preferentially retained at intracellular membranes (this study and (18, 19)). We propose that the hERG1a PAS domain-N-linker region stabilizes hERG1b through a mechanism where the PAS domain of hERG1a interacts directly with the C-linker and CNBHD of hERG1b (this study and (16)) and the N-linker region of hERG1a interacts with the N-terminal domain of hERG1b (Fig. 9A). We propose that residues ^216^TAMDN^220^ in the hERG1a N-linker are required for the functional upregulation of hERG1b, perhaps by being necessary for the interaction of the N-linker with hERG1b. In heteromeric channels comprising full-length hERG1a and hERG1b subunits, such as those in the heart (1, 2), we propose that the intracellular N-terminal domains are arranged such that the PAS domain of hERG1a makes an intersubunit interaction with the CNBHD of hERG1b and that the hERG1a N-linker domain interacts with the hERG1b N-terminal domain (Fig. 9B). These interactions help to promote the surface expression of hERG1b in the presence of hERG1a.

**Figure 9.**
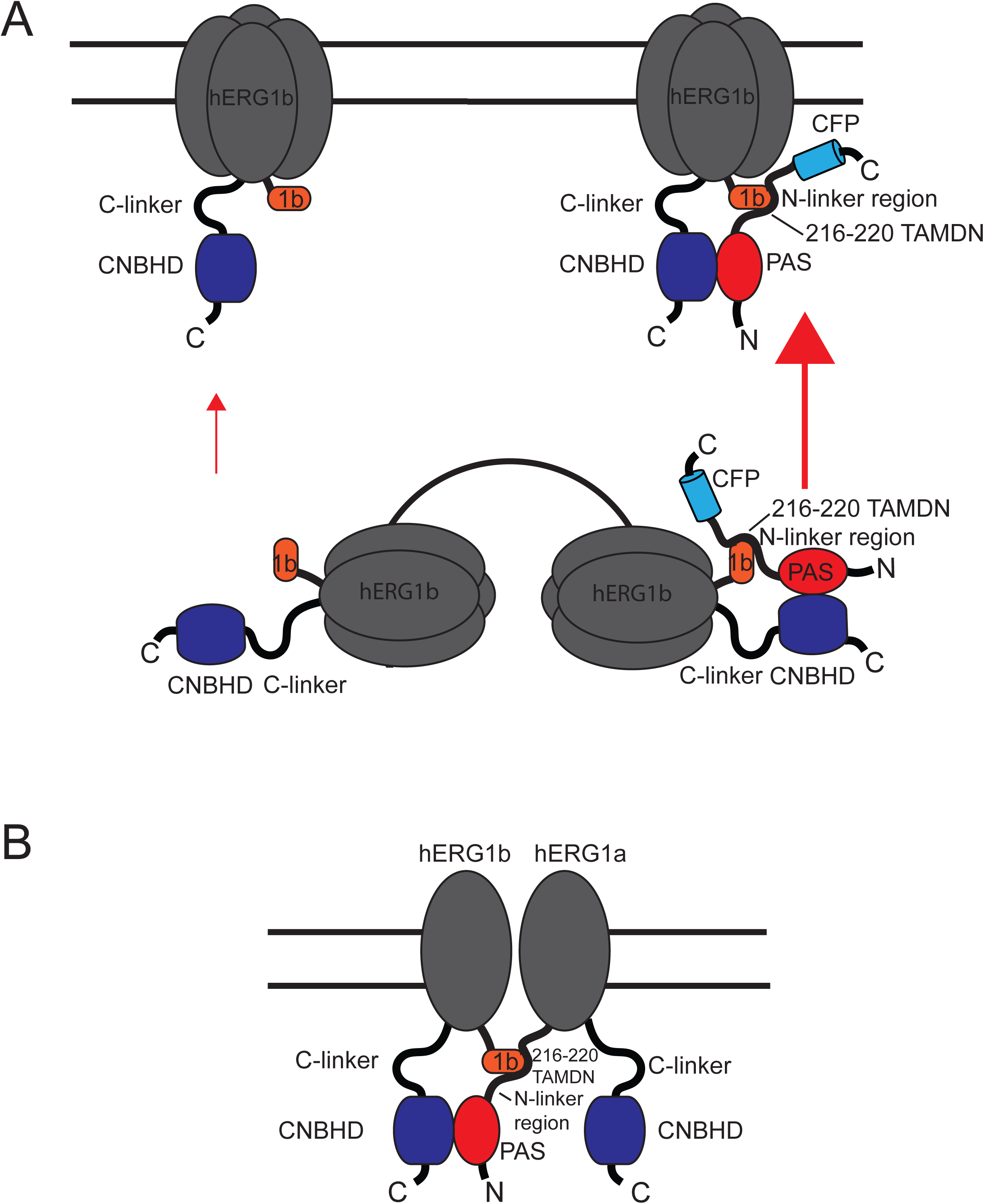
Scheme of hERG1a PAS-N-linker domain upregulation of hERG1b at the plasma membrane. We propose that A) the hERG1a PAS domain interacts with the CNBHD of hERG1b and that the hERG1a N-linker interacts with the hERG1b domain and increases (large arrow) hERG1b channels at the plasma membrane, relative to wild-type hERG1b channels (small arrow). B) In heteromeric hERG1a/hERG1b channels, we propose that the N-terminal domains of hERG1a make an intersubunit interaction with hERG1b subunits.

Our results here fit well with previous conclusions about hERG1a and hERG1b N-terminal domain interactions and the location of a region in hERG1a that is necessary for upregulation of hERG1b. In a previous study, a biochemical interaction between fusion proteins encoding the hERG1a N-terminal domain-S2 transmembrane domain and hERG1b N-terminal domain-S2 transmembrane domain was detected (18). Here, we refine the identity of the interacting regions by showing that the hERG1a PAS domain-N-linker and the hERG1b N-terminal domain were sufficient to form an interaction (Figs. 3,4). In a previous study, full-length hERG1a and hERG1a subunits with a deletion of amino acids 2-200 both increased the maturation of hERG1b, indicating that an enhancer region for hERG1b was upstream of reside 200 (19). In this study, we mapped the region necessary for hERG1a to upregulate hERG1b to amino acids 216-220 in the hERG1a N-linker (Figs. 6-8).

Di-arginine motifs in the hERG1b unique N-terminal region are important for ER retention of hERG1b and are masked by hERG1a, leading to an increase in hERG1b at the plasma membrane (19), which we confirm here (Fig. S2). We tested whether hERG1b di-arginine motifs were required for the increase in hERG1b by the hERG1a PAS domain-N-linker region. We report that a hERG1a PAS-N-linker region (hERG1a N1-228) increased hERG1b channels bearing di-arginine motif mutants at the plasma membrane as measured with biotinylation, suggesting that the hERG1a PAS domain-N-linker region-dependent increase in hERG1b is by a mechanism independent of hERG1b di-arginine motifs (Fig. S2). We speculate that another part of hERG1a, besides the N-terminal PAS-N-linker domain, masks hERG1b di-arginine motifs.

We detected a larger leftward shift in the voltage dependence of activation of hERG1b Δ2-36 co-expressed *in trans* with the hERG1a PAS domain-N-linker region than hERG1b co-expressed *in trans* with hERG1a PAS domain-N-linker region (Fig. 2A, C, Table 1). This was unexpected as the 1b domain has been previously associated with housing the RXR motif in trafficking, but not as a determinant of gating. The mechanism for this leftward shift in not clear, but it is consistent with an additional role of the hERG1b 1b domain in hindering activation of hERG1b by the hERG1a PAS domain-N-linker.

The direct interaction of mammalian ERG1a and ERG1b is crucial for cardiac physiology. Mammalian ERG1a and ERG1b subunits interact to form native cardiac I^Kr^ in the mammalian heart, including human heart, and manipulation of I_Kr_ to favor hERG1b-like kinetics or hERG1a-like kinetics is proarrhythmic (1-3, 8, 18-22). Our results indicate that the N-linker of hERG1a is a new player in this assembly mechanism and thus disruption of the N-linker mechanism may be a target for inherited Long QT syndrome mutations and a target for therapeutic interventions.

## Acknowledgements

Research reported in this publication was supported by the National Institutes of Health/National Institute of General Medical Sciences R01 GM130701 and R01 GM127523 (MCT) and the Training Program in Integrative Membrane Biology, T32 008181 (TRC) and the National Heart, Lung and Blood Institute F32HL131189-01 (AAJ).The authors thank Dr. Beth McNally for assistance with experiments. We thank Dr. Gail A. Robertson for sharing hERG1b RXR mutant channels.

## Materials and Methods

### Molecular Biology

The hERG1a and hERG1b constructs used here were described previously (16, 24). Some recombinant hERG1a N-terminal domains were described previously (16, 24) but hERG1a N1-200-mCFP and hERG1aN1-228-mCFP with alanine mutants, hERG GST fusion proteins and 6xHis/Flag-tagged fusion proteins were made with customized primers and generated commercially (BioInnovatise, Inc. Rockville, MD). CFPs were included to aid in visual detection, detection on Western blots and as FRET donors. Constructs used in FRET 2-hybrid studies were either previously described (12) or generated commercially (i.e. hERGb-1-56-Citrine, hERG1a 136-354-mCFP or hERG1a 666-1159-Citrine) (Bioinnovatise). For oocyte expression, all constructs were in the pGH19 vector and for HEK293 cell expression all constructs were in the pcDNA3.1 vector. RNA was transcribed in vitro with the T7 mMESSAGE mMACHINE kit (Invitrogen).

### Electrophysiology

Two-electrode voltage-clamp and whole-cell patch-clamp studies were performed as previously described ((24, 25). Oocytes were obtained from a commercial source (Ecocyte, Inc.) and injected with 49 nl of RNA and incubated for 24-72 hours in ND-96 buffer + 50 μg/mL gentamycin prior to electrophysiology. A ratio of 2:1 for N-terminal domains to channel RNA was used in co-expression studies. TEVC was performed using (OC-725C, Warner Inst.) and an A/D converter (ITC-18, Instrutech). Recording electrodes were fashioned with a micropipette puller (Sutter), filled with 3M KCl and had resistances of 0.5 to 1.2 MOhm. The external (bath) solution was (mM) 4KCl, 94 NaCl, 1 MgCl2, 0.3 CaCl2, 5 HEPES with pH 7.4.

HEK293 cells were transiently transfected with TransIT-LT1 reagent (Mirus) with channel cDNA and hERG1a N-terminal domain cDNAs in a 2:1 ratio in co-expression experiments. Whole cell recordings were performed 24-48 hours after transfection using a patch-clamp amplifier with a built-in A/D converter (EPC10, HEKA, Inc.). Microelectrodes had resistances of 2-3 MOhm after filling with pipette (internal) solution of (in mM) 130 KCl, 1 MgCl2, 5 EGTA, 5 MgATP and 10 HEPES, pH 7.2. The bath (external) solution was (in mM) 137 NaCl, 4 KCl, 1.8 CaCl2, 1 MgCl2, 10 glucose, 10 HEPES, pH 7.4. Capacitance was compensated on-line. No leak subtraction was performed.

Electrophysiological recordings were made at room temperature and acquired with Patchmaster Data Acquisition software (HEKA Electronic). Currents were analyzed off-line using Igor Pro (WaveMetrics). Conductance-voltage data was fit with a Boltzmann function (y = 1/ [1 + e[(V_1/2_ - V)/k]]) in which V_1/2_ is the half-maximum potential for activation and k is the slope factor. Deactivating currents were fit with a single exponential function. (y = Ae(–t/tau)) where t is time and tau is the time constant of deactivation. All data are presented as the mean ± SEM. *n* is the number of cells. One-way ANOVA with Tukey’s test was performed to determine statistical significance. A value of P < 0.01 was considered statistically significant.

### Biochemistry and biotinylation

Biochemical detection of hERG1a and hERG1b proteins was performed using biotinylation as previously described (24). Briefly, HEK293T cells were transfected with hERG1b or hERG1a cDNAs using TransIT-LT1 transfection reagent (Mirus). After 24-48 hours cells were incubated with extracellular application of EZ-link Sulfo-NHS-SS-biotin (1mg/mL, Pierce) and proteins were purified with biotin-streptavidin beads. Bead-bound proteins were separated by SDS-PAGE, transferred to nitrocellulose and immunoblotted with a primary antibody (anti-hERG KA, ENZO; anti-GFP ab290, Abcam) and an HRP-conjugated secondary antibody (goat anti-rabbit, Life Technologies). Blots were visualized with a gel imager (BioRad XRS) using chemiluminescence (SuperSignal West Femto, ThermoFisher).

### Biochemical pull-down interaction assay

Biochemical interaction assays were performed as previously described (13). Briefly, Glutathione *S*-transferase (GST) hERG1a N-terminal domains or GST-only negative controls constructs were grown in BL21 (DE3) competent cells (Agilent Technologies) until they reached exponential growth. Protein expression was induced using isopropyl-β-d-thiogalactoside (0.4-mM; Research Products International) with shaking at 30 degrees Celsius overnight. Bacteria were harvested with centrifugation, resuspended in buffer S (50-mM Tris, pH 8, 150-mM NaCl, 25-mM imidazole, 0.5% CHAPS, and 0.25% Tween 20; Sigma-Aldrich) and lysed with a sonicator. The products were centrifuged at high speed and the supernatants were applied to glutathione beads (GE Healthcare), washed with buffer S and the protein concentration was measured using a Bradford assay.

6xHis fusion proteins were grown in M15 cells (VWR) and prepared as above except that after IPTG induction, proteins were grown O.N at 18 degrees C. Materials were cleared by a highspeed spin as above and applied to a nickel column (HiTrap Chelating HP, GE Healthcare), washed and eluted in buffer S with imidazole (500 mM).

To perform interaction assays, the GST fusions were bound to beads and 6xHis fusion proteins were combined in Buffer S with 0.5 % CHAPS detergent. Proteins were incubated O.N. at 4 degrees Celsius on a rotating mixer. Samples were washed with Buffer S, and proteins were stripped from beads by incubation in gel loading buffer with beta-mercaptoethanol and loaded to 4-15 % gels (Criterion; BioRad) for SDS-PAGE. GST inputs were assessed by Coomassie Blue staining of gels. Interacting proteins were resolved with SDS-PAGE, transferred to nitrocellulose and blotted with an anti-Flag antibody (Sigma-Aldrich) and HRP-linked secondary antibody (ThemoFisher) and visualized with a chemiluminesce detection kit (SuperSignal West Femto; ThermoFisher) and a gel imaging system (ChemiDoc XRS, BioRad).

### FRET two-hybrid assay

A FRET hybridization assay was used to measure an association between two proteins labelled with a donor (mCFP) or acceptor (Citrine). As in previous work, we used a spectral separation method for analysis (24), using isolated domains of channels fused to CFP or Citrine (12, 23). Spectra were measured from HEK293 cells using a 60x objective with NA 1.45 (Nikon) and an inverted microscope (Nikon TE-2000). The excitation source was a 120 W lamp (X-Cite 120), and emission was detected with a spectragraph (SpectraPro 2150i; Acton Research) and a CCD camera (Roper 512B, Roper Scientific). MetaMorph 6.3r7 (Universal Imaging/Molecular Devices) and Monochromator Control (Acton) acquisition software packages were used to take images and spectrum. To measure fluorescence, a “FRET” cube with a long pass emission filter (D436/20, 455dclp, D460lp; Chroma) was used to obtain CFP only traces (cyan traces Fig. 4) or traces from cells expressing CFP and Citrine-containing fusion proteins (blue traces, Fig. 4). The acceptor fluorescence was measured with a YFP cube (green trace, Fig. 4). An extracted spectrum (red trace, Fig. 4) was generated that contained the direct excitation of Citrine, F_direct436_, and the excitation of Citrine due to FRET, F_FRET436_, by subtracting a scaled CFP-only control spectrum (cyan trace). The resulting ratio (ratio of red/green trace) is Ratio A = F_direct436_ + F_FRET436_ /F_500_ which contains a direct component and FRET component. RatioA_0_ was also generated which is the red trace/green trace (F_direct436_/F_500_) in a control experiment with hERG1a Citrine (not shown). Ratio A - RatioA_0_ is equal to F_FRET436_/F_500_ which is a value proportional to FRET efficiency (26, 27).

