## Supplemental Figures 1 and 2 for "The N-linker region of hERG1a upregulates hERG1b potassium channels"

Figure S1

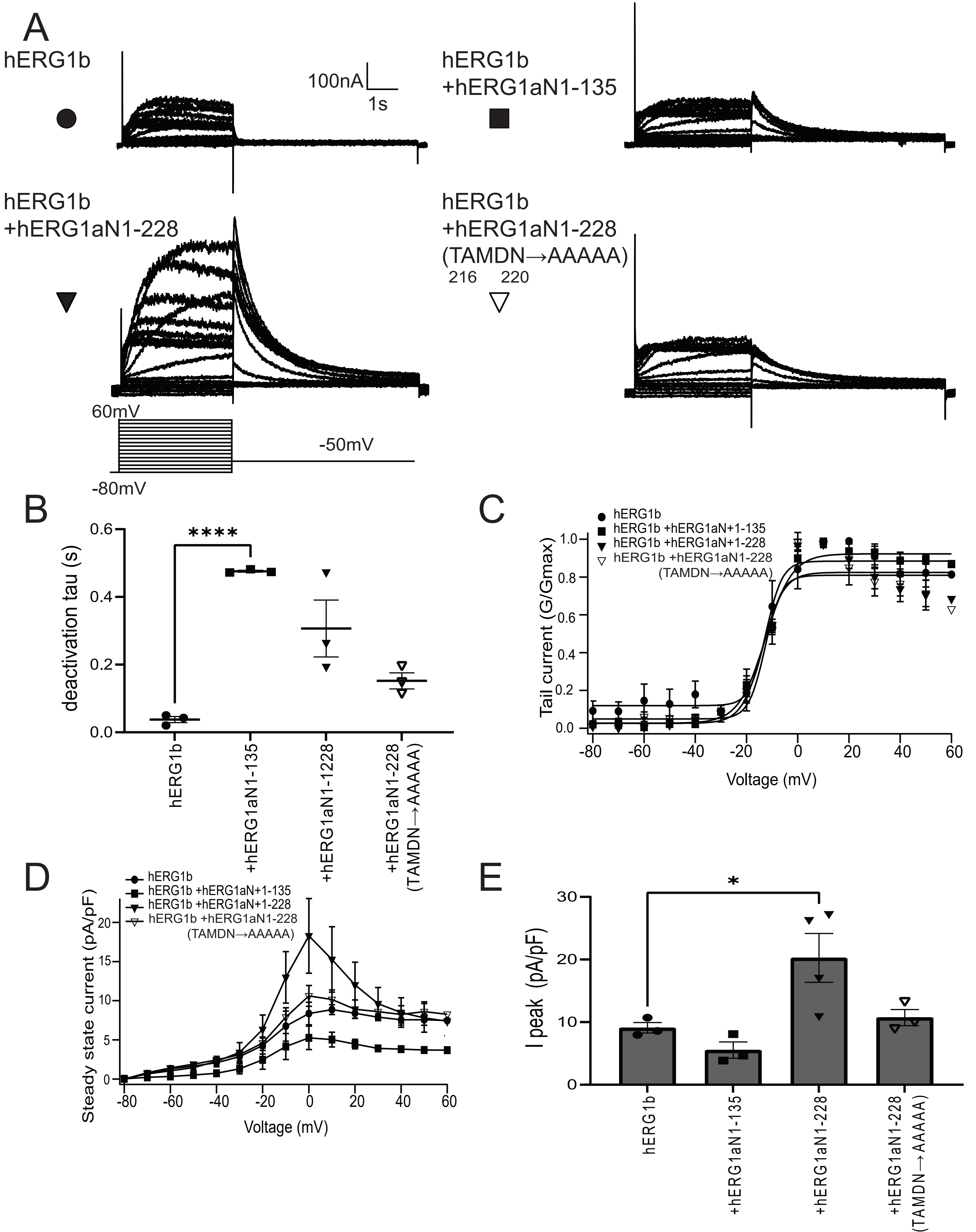
A
hERG1b
100nA
1s
hERG1b
+hERG1aN1-135
hERG1b
+hERG1aN1-228
hERG1b
+hERG1aN1-228
(TAMDN→AAAAA)
216 220
60mV
-50mV
-80mV
B
deactivation tau (s)
****
hERG1b
+hERG1aN1-135
+hERG1aN1-1228
+hERG1aN1-228 (TAMDN→AAAAA)
C
hERG1b
hERG1b +hERG1aN+1-135
hERG1b +hERG1aN+1-228
hERG1b +hERG1aN1-228 (TAMDN→AAAAA)
Tail current (G/Gmax)
Voltage (mV)
D
hERG1b
hERG1b +hERG1aN+1-135
hERG1b +hERG1aN+1-228
hERG1b +hERG1aN1-228 (TAMDN→AAAAA)
Steady state current (pA/pF)
Voltage (mV)
E
*
I peak (pA/pF)
hERG1b
+hERG1aN1-135
+hERG1aN1-228
+hERG1aN1-228 (TAMDN→AAAAA)

Figure S2

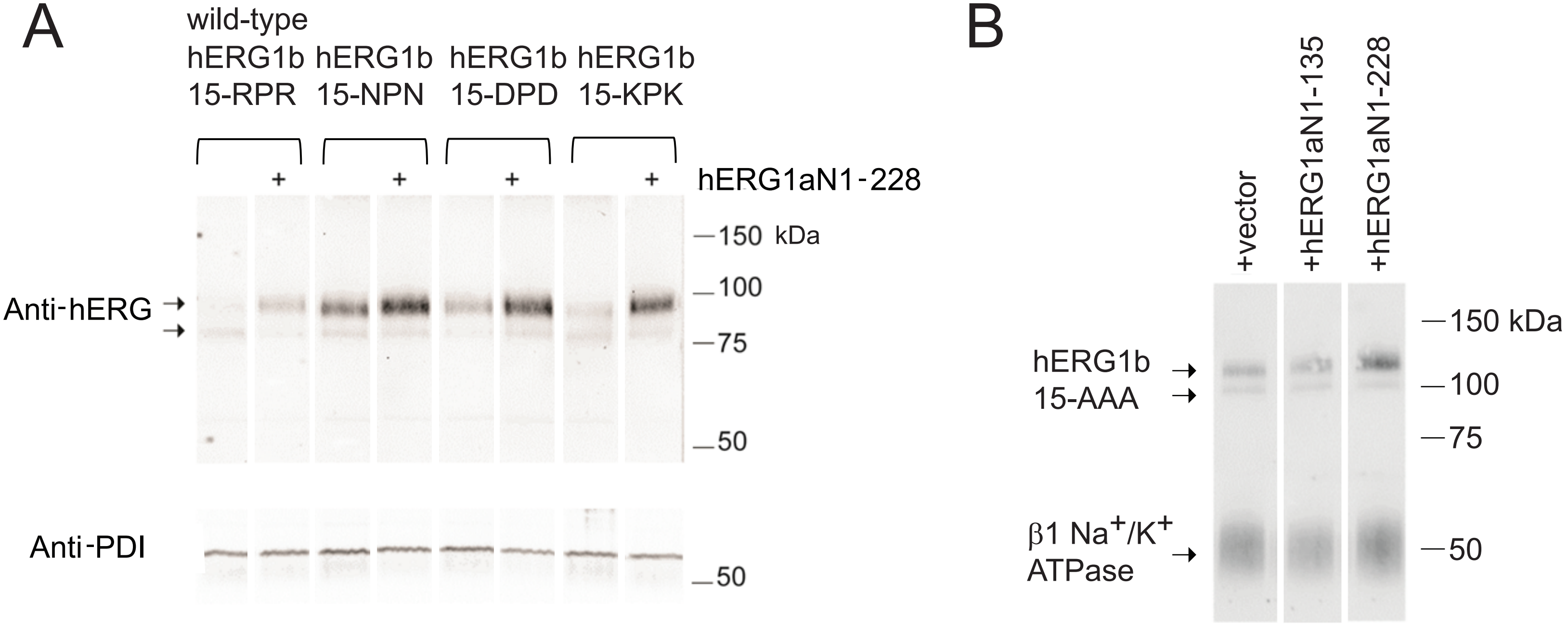

A
wild-type
hERG1b hERG1b hERG1b hERG1b
15-RPR 15-NPN 15-DPD 15-KPK
+ + + +
hERG1aN1-228
150 kDa
100
75
50
Anti-hERG
Anti-PDI
50
B
+vector
+hERG1aN1-135
+hERG1aN1-228
150 kDa
100
75
50
hERG1b
15-AAA
β1 Na+/K+
ATPase

**Figure S1. hERG1a N-terminal PAS and N-linker domain increases hERG1b currents in HEK293 cells.** A) Whole-cell patch-clamp recordings of hERG1b, hERG1b + hERG1a PAS domain (hERG1a N1-135), hERG1b + hERG1a PAS and N-linker domain (hERG1a N1-228), and hERG1b + hERG1a N1-228 $^{216}$TAMDN$^{220}$ mutated to $^{216}$AAAAA$^{220}$. B) Time constant of deactivation from fit to tail currents at -50mV. C) Conductance-voltage (G-V) plot. D) Current-voltage (I-V) plot. E) Histogram of peak current amplitude at depolarizing voltage. N ≥ 3 for each. Error bars are mean ± SEM.

**Figure S2. hERG1a PAS domain -N-linker regions enhance surface expression of hERG1b channels with mutations in the di-arginine (RXR) ER retention motif** A) Western blot of biotinylated hERG1b proteins as indicated, hERG1b 15-RPR (wild-type hERG1b with RXR motif at amino acids 15-17), hERG1b 15-NPN, hERG1b 15-DPD and hERG1b 15-KPK expressed alone or *in trans* (+) with the hERG1a PAS domain-N-linker (hERG1aN1-228). The loading control was PDI. In this experiment, hERG1b and hERG1b RXR mutants (a kind gift from Dr. G.A. Robertson) were not labelled with Citrine. B) Western blot of biotinylated hERG1b-Citrine with 3 alanine mutations at the RXR motif (RXR to AAA) expressed with empty vector, the hERG1a PAS domain (N1-135) and the hERG1a PAS domain-N-linker region (hERG1aN1-228). Loading control is Na/K ATPase.
